# Unraveling the Regenerative Proteomic Signature of *Helix aspersa*’s Slime in Human Dermal Fibroblasts by Data-driven Proteomics Approach

**DOI:** 10.64898/2026.03.08.709924

**Authors:** Muhammad Rashad, Alessia Ricci, Marwa Balaha, Zsuzsanna Darula, Ádám Pap, Amelia Cataldi, Éva Csősz, Susi Zara

## Abstract

Proteins orchestrate essential cellular processes, including metabolism, communication, survival, and regeneration, making proteomic profiling a powerful strategy to elucidate complex biological responses. Snail slime (SnS) has emerged as a bioactive material with documented pro-healing, antioxidant, and anti-inflammatory properties; however, its effects at the proteome level in normal human dermal fibroblasts (NHDFs) remain unexplored. In this study, an LC–MS-based proteomic approach (Data are available via ProteomeXchange with identifier PXD075292) combined with network and Gene Ontology enrichment analyses was employed to investigate SnS-induced molecular reprogramming in NHDFs, followed by functional assays. Results show that SnS is well tolerated for up to 72 h, confirming its cytocompatibility, followed by proteomic analysis revealing enrichment of biological processes related to apoptosis regulation, oxidative stress response, wound healing, cell migration, and anti-aging. Network analysis identified AKT, PI3K, SRC, and KRAS family members as key hub proteins, indicating convergence on central signaling pathways controlling survival, redox balance, and migratory activity. Functional assays demonstrated a time-dependent, controlled modulation of apoptosis consistent with cellular turnover, alongside a hormetic redox response characterized by transient ROS signaling followed by enhanced antioxidant capacity. Importantly, SnS significantly accelerated fibroblast migration, achieving complete wound closure within 24 h. Collectively, these findings demonstrate that SnS induces coordinated proteomic and functional reprogramming that integrates redox modulation, controlled apoptosis, and enhanced migration, providing a mechanistic basis for its pro-healing and anti-aging effects and supporting its potential as a regenerative biomaterial.

## 1. Introduction

*Helix aspersa*, more accurately classified as *Cornu aspersum*, is a terrestrial gastropod mollusk widely distributed across Europe, North Africa, and other temperate regions due to both natural dispersion and human-mediated introduction [1]. Belonging to the phylum Mollusca, class Gastropoda, and family Helicidae, this species is commonly referred as the “garden snail” and is among the most studied land snails due to its ecological adaptability and economic importance [2]. Its biological characteristics, particularly its capacity to secrete mucus, have attracted growing scientific interest for applications extending far beyond its role in agriculture or cuisine [3].

Snail slime (SnS), or mucus, is a complex biological secretion that plays a vital role in the survival and locomotion of *H. aspersa* [4]. Originally understood as a means to aid movement across rough terrain, prevent desiccation, and deter predators, this viscous substance has revealed a remarkable biochemical composition with significant biomedical and cosmetic potential [5]. The mucus is produced primarily by specialized glands in the foot and mantle of the snail [6] and contains a diverse array of compounds, including glycoproteins, enzymes, antimicrobial peptides, allantoin, collagen, elastin, and glycolic acid [6–8]. These components are essential not only for the snail’s own protection and wound healing but also hold promise for human applications, especially in the fields of dermatology, regenerative medicine, and pharmaceuticals [9].

Interest in *H. aspersa* mucus dates back to traditional medicine practices, particularly in Mediterranean regions, where snail-based remedies were used for skin inflammation and other ailments [4]. In recent decades, advances in molecular biology and biochemistry have facilitated the identification and characterization of the active constituents in snail slime, enabling its incorporation into modern skin care formulations and experimental therapeutics [10]. As a result, what was once a naturally evolved adaptation for terrestrial life has emerged as a valuable bioresource with increasing relevance in health and biomedical sciences.

Advancements in the extraction of SnS have also enabled its production and use in versatile fields, not limited to cosmetics only. Advanced cruelty-free extraction Muller method, for example, provides the basis for meeting the market expectations without endangering the species. It helps to obtain SnS without compromising the product quality, while preserving the animals’ life [4]. The optimum quality SnS containing a significant amount of bioactive macromolecules [6], when applied to human skin, promotes dermal fibroblast viability [10, 11], exhibits antioxidant activity [12, 13], modulates the release of reactive oxygen species (ROS) [14–16], counteracts inflammation [10], and has the ability to stimulate cell proliferation and migration [17].

Proteomics refers to the comprehensive analysis of the structural and functional proteins expressed in a biological samples, providing insights into the physiological state of an organism at a specific point in time [18]. In contrast to the relatively static nature of the genome, the proteome is highly dynamic, reflecting ongoing biological processes, including responses to environmental factors, pathological conditions, and therapeutic interventions [19]. A study conducted by Lopes *et al.* on sea slug *Elysia crispate* mucus using HPLC/MS/MS identified 306 abundant proteins with a majority of hydrolases accounting for 47% of all enzymes, suggesting antimicrobial activity [20]. Similarly, Li *et al.* hydrolyzed the *Helix aspersa* slime with trypsin and analyzed by LC-MS to obtain active peptides such as EK-12 (mw: 1366.2 Da) [21]. Furthermore, Cerullo *et al.* applied LC-MS/MS based proteomic approach to purified mucus obtained from *Cornu aspersum* using shotgun proteomics and detected 71 proteins [22]. However, to date, proteomic investigations have primarily focused on raw or purified snail mucus, often analyzed directly or following gel electrophoresis separation. While these studies have provided valuable information on mucus composition and identified abundant proteins and peptides, they offer limited insight into the biological effects of SnS on human cells. This gap highlights the need for MS-based proteomic approaches not only to address the analytical challenges posed by viscous and complex biological matrices but also to elucidate the functional cellular responses elicited by SnS exposure.

In our previous studies, we tested the biological effect of SnS on different cell types, including human keratinocytes (HaCaT), gingival and dermal fibroblasts (HGFs and NHDFs, respectively), human macrophages, and endothelial cells (ECs) [10, 15, 23]. Being the major population of skin, NHDFs were chosen for further exploration of proteomics profiling after SnS treatment.

While previous studies investigating the biological effects of SnS on fibroblasts have primarily focused on individual proteins, typically analyzed via conventional techniques such as Western blotting and enzyme-linked immunosorbent assay (ELISA), a comprehensive proteomic characterization has remained lacking. To address this gap, the present study not only confirmed the previously reported effects of SnS on cell viability and its antioxidant properties, including ROS downregulation, but also employed a proteomics-based approach to systematically analyze the protein expression profile of SnS-treated NHDFs. High-resolution EvoSep ONE liquid chromatography coupled with ultra-sensitive Orbitrap mass spectrometry was employed to achieve an in-depth characterization of proteomic alterations induced by SnS treatment.

## 2. Materials and methods

### 2.1 SnS extraction

The SnS was extracted by using the cruelty-free Cherasco Muller method, described in detail in our previous studies [4, 6, 10]. The extracted SnS was then filtered through a 0.2 micron filter and mixed with stabilizers (0.1% of sodium benzoate and potassium sorbate) to enhance quality and productivity [23].

### 2.2 Cell culture and treatment

NHDFs, purchased from Merk Life Science, Milan, Italy (catalog no. c-12302, Lot#505Z033.1), were cultured in high-glucose Dulbecco’s Modified Eagle’s Medium (DMEM) supplemented with 1% penicillin/streptomycin antibiotics and 10% fetal bovine serum (FBS) at 37℃ with 5% CO_2_. Cells were treated with three different dilutions of SnS 1:40 (0.508 mg/mL), 1:60 (0.31 mg/mL), and 1:80 (0.247 mg/mL), established previously, and the control was treated with stabilizers only. All the treatments were prepared in FBS-free DMEM to obtain the effect of SnS solely on various biological functions evaluated in this study.

### 2.3 Cell viability assay

Cell viability was assessed using the Alamar Blue assay, with all experimental conditions run in triplicate. NHDFs were plated at a density of 6700 per well in a 96-well plate. After 24, 48, and 72 h of culture, the medium was replaced with fresh medium containing 10% (v/v) Alamar Blue reagent (Thermo Scientific, Rockford, IL, USA). The plates were then incubated for 4 hours at 37 °C. Following the incubation, the absorbance was measured at 570 nm and 600 nm. A well containing only culture medium and reagent, without cells, served as the negative control. The percentage reduction of the Alamar Blue reagent was determined according to the manufacturer’s protocol.

### 2.4 Treatment and sample collection for proteomic analysis

NHDFs were grown in culture-treated T75cm^2^ flask to reach 80-90% confluency, washed twice with Dulbecco’s Phosphate Buffered Saline (PBS), and then treated with 1:40, 1:60 and 1:80 dilutions of SnS for 24 and 48 h and control for 0, 24 and 48 h in FBS-free DMEM in triplicate. After the aforementioned time points, the medium was removed, and cells were collected by trypsinization, washed with DPBS and pellets were stored in -80 ℃ for future use.

### 2.5 Cell lysis and protein quantification

All the SnS treatments were performed in triplicate, and the collected samples were grouped in order to avoid handling errors between the biological replicates. The samples were divided into three equal groups by randomly selecting one sample from each treatment condition (3 x 9 samples). Samples in a group/block were processed together. Each cell pellet was lysed with 100 μL of S-TRAP 1X lysis buffer (5% sodium dodecyl sulfate (SDS) + 50 mM triethylammonium bicarbonate buffer (TEAB), vortexed, and subjected to probe sonication (HD 2200, Bandelin Sonopuls, Berlin, Germany) for 40 sec (4 cycles x10%) at 50% power (MS 72). Samples were centrifuged at 13000 xg for 8 min, and the supernatant was collected. The protein amount in each sample was measured by the bicinchoninic acid assay (BCA) (Merk Life Science, Milan, Italy).

### 2.6 Protein digestion by S-TRAP protocol

100 μg protein from each sample was digested using the S-TRAP protocol, as previously described by Zougman *et al.* [24]. Proteins were reduced by incubating for 15 min at 55 °C with 10 mM dithiothreitol (DTT) final concentration, followed by sulfhydryl group alkylation by incubating for 10 min at room temperature with 20 mM iodoacetamide (IAA) final concentration. Samples were acidified with 12% phosphoric acid to a final concentration of 1.2%. Each sample was diluted 6X with binding buffer (100 mM TEAB in 90% methanol) and filtered through S-TRAP mini spin column (ProtiFi, Fairport, NY, USA) at 4000 xg for 30 sec to trap proteins. Samples were washed three times (at 4000 xg for 30 sec) with binding buffer then incubated overnight at 37 °C with trypsin/Lys-C mix (Pierce Trypsin/Lys-C Protease Mix, MS Grade, USA, Lot# YJ380698) in 50 mM TEAB at a 1:25 enzyme to protein ratio (w/w). Peptides were eluted with 80 μL of 50 mM TEAB, 0.2% FA (formic acid), and 50% ACN (acetonitrile) at 4000 xg for 1 min stepwise and dried in a speedvac (RVT 400, Thermo Scientific, USA).

### 2.7 LC-MS analysis

Samples were redissolved in 0.1% FA and sample aliquots representing 1 μg protein digests were analyzed using an EvoSep ONE liquid chromatography (LC) (Evosep Biosystems, Odense, Denmark) coupled to an Orbitrap Fusion Lumos Tribrid (ThermoFisher Scientific, Waltham, Massachusetts, USA) mass spectrometer (MS). The analytical column, EvoSep Endurance column (EV1106) (Evosep Biosystems, Odense, Denmark): 150 mm (L) × 0.15 mm (D), ReproSil-Pur C18, 1.9 µm beads, with a column temperature of 30 °C, was used for peptide fractionation. A 44-minute gradient program, 30 samples per day (SPD) was employed using two solvents (Solvent-A: 0.1% FA in water, Solvent-B: 0.1% FA in ACN). Samples were measured using data-independent acquisition (DIA using a 50 %-staggered 12 Da isolation windows scheme. A complete MS data acquisition method is described in Supplementary Table S1. The mass spectrometry proteomics data have been deposited to the ProteomeXchange Consortium via the PRIDE [25] partner repository with the dataset identifier PXD075292.

### 2.8 Data analysis

The raw files for each sample’s analysis were run in DIA-NN (v. 2.2) against a human FASTA file having 20431 entries, downloaded from www.uniprot.org. Missed cleavages were set to 2 and a maximum of 3 variable modifications along with protein N-terminal M-excision, carbamidomethylation of cysteine, oxidation of methionine and protein N-terminal acetylation. Peptide length was set to 5-30, precursor charge range 1-4, precursor m/z range 300-1800 and fragment ion m/z range 200-1800 with active match between runs (MBR). The DIA-NN output file was then used to sort the protein groups (PG) with at least 2-peptide hits with the help of the in-house developed KNIME analytics platform (manuscript under consideration in Journal of Proteomics Research).

Differentially abundant proteins (DAPs) were extracted by considering the unique proteins in a specific treatment group plus the proteins with a statistically significant change between the groups. The data was refined and sorted with the help of RStudio software (v. 2025.05.01, packages: “readr”, “dplyr”, and “purrr”), and statistical significance value was calculated by using GraphPad Prism v.8.0 software (San Diego, CA, USA), after applying Multiple t-test followed by Benjamini, Krieger, and Yekutieli method FDR correction (q<0.05). The unique and common proteins were visualized by Venn diagram created with Venny 2.1.0. The network analysis was performed by using string-db.org (v.12.0) [26] with stringency set to the highest (0.9). The DAPs plus their maximum 50 first interactors were considered, and disconnected nodes were kept hidden. Functional enrichment analysis was carried out in String, merging the GO terms showing a similarity higher than 0.8. The selected terms were used for visualization in String. String networks were exported to Cytoscape v3.10.4. [27] and the hub proteins were examined with CytoHubba plugin.

### 2.9 Antioxidant activity

The total antioxidant capacity (TAC) of SnS was determined by evaluating its power to reduce Cu²⁺ to Cu⁺ using the antioxidant assay kit (catalog number MAK334, Merck Life Science, Milan, Italy) according to the manufacturer’s instructions. NHDFs were seeded in triplicate in a 6-well plate at a cell density of 200,000 cells/well. After 24 h of SnS treatment (1:40, 1:60, 1:80), cell supernatants were collected for each condition to measure TAC in cell lysate. Briefly, 20 µL of cell supernatants were pleated in a flat bottom 96-well plate and 100 µL of Reaction Mix were added and maintained for 10 min at room temperature. Absorbance was read at 570 nm wavelength by using a spectrophotometer (Multiskan GO, Thermo Scientific, Waltham, MA, USA) and values were interpolated with a standard Trolox curve to obtain the concentration of antioxidant molecules (µM).

### 2.10 Flow cytometry analysis of intracellular ROS production

NHDFs were seeded in triplicate in 6-well plates at a density of 180,000 per well and exposed to SnS at dilutions of 1:40, 1:60, and 1:80 for 6 and 24 h. Intracellular ROS release was measured after 6 and 24 h of SnS treatment rather than 24 and 48 h, primarily due to the kinetics of ROS generation and the short half-life of these molecules. After treatment, cells were incubated with 5-(and-6)-chloromethyl-2′,7′-dichlorodihydrofluorescein diacetate (CM-H₂DCFDA, Cat. C6827, Molecular Probes, Invitrogen, Life Sciences Division, Milan, Italy; final concentration of 5 μM) for 1 h at 37 °C, following the procedure described previously [28]. Intracellular ROS production was detected as an increase in green fluorescence resulting from probe oxidation and measured using a CytoFLEX flow cytometer (Beckman Coulter, FL, USA) equipped with an FL1 detector and operated in logarithmic mode. Data acquisition and analysis were performed using the CytExpert software, and ROS production was quantified as the median fluorescence intensity (MFI) ratio obtained from histogram statistics. Dead cells were excluded by staining with propidium iodide (PI; 5 μg/mL, Cat. P4864, Merk Life Science, Milan, Italy). For each sample, a minimum of 10,000 events were acquired for analysis.

### 2.11 Detection of Apoptosis and Necrosis by Flow Cytometry

NHDFs were seeded in triplicate in 6-well plates at 200,000/well, and following 24 and 48 h treatments with SnS (1:40, 1:60, and 1:80 dilutions), both cells and their corresponding supernatants were collected. To distinguish apoptotic and necrotic cell populations, the cells were stained using an Annexin V and Propidium Iodide (PI) kit (eBioscience, Thermo Fisher Scientific), following the manufacturer’s protocol. Briefly, cells were incubated in the dark for 15 min at room temperature in a binding buffer solution containing Annexin V-FITC and PI. Fluorescence quantification was performed on a CytoFlex flow cytometer (Beckman Coulter). The FITC signal from Annexin V was detected using a 530/30 bandpass filter (FL-1 channel), while PI fluorescence was measured with a 650 nm long pass filter (FL-3 channel). For each sample, around 10,000 events were acquired, and the data were processed using CytExpert 5.0 Software (Beckman Coulter). Cell populations were quantified based on their staining profiles on density plots: viable cells (Annexin V⁻/PI⁻) were identified in the lower left quadrant; early apoptotic cells (Annexin V⁺/PI⁻) in the lower right quadrant; late apoptotic cells (Annexin V⁺/PI⁺) in the upper right quadrant; and necrotic cells (Annexin V⁻/PI⁺) in the upper left quadrant.

### 2.12 Western Blot Analysis

Cells were seeded in triplicate in 6-well plates (200,000 cells/well) and following 24 and 48 h treatments with SnS (dilutions 1:40, 1:60, 1:80) were evaluated. After treatment, cells were detached with trypsinization and centrifuged at 1200 rpm for 10 min at 4°C to obtain a pellet. The pellets were lysed using RIPA buffer supplemented with a protease inhibitor cocktail (sodium orthovanadate, leupeptin, aprotinin, PMSF; Merck Life Science, Milan, Italy). The lysates were then centrifuged at 15,000 x g for 15 min at 4°C, and the resulting supernatants were collected for protein quantification via the BCA method (QuantiPro™ BCA Assay kit, Merck Life Science, Milan, Italy), adhering to the kit’s protocol. For immunoblotting, 30 µg of total protein per sample was resolved on 10 % SDS-polyacrylamide gels and subsequently transferred to nitrocellulose membrane using Invitrogen Power Blotter (ThermoFisher Scientific, Waltham, Massachusetts, USA). Membrane blocking was performed with 5% non-fat milk. The blots were then probed overnight at 4°C with the following primary antibodies: anti-Bcl-2 (catalog no. sc-7382, mouse, 1:100), and anti-Bax (catalog no. sc-7480, mouse, 1:100) purchased from Santa Cruz Biotechnology (Dallas, Texas, USA), and anti-Actin (catalog no. MA1-80729, mouse, 1:10000) purchased from ThermoFisher Scientific (Waltham, Massachusetts, USA). Following primary incubation and washes with PBS-Tween, membranes were exposed to appropriate HRP-conjugated secondary antibodies (Calbiochem, Merck Life Science, Milan, Italy). Signal detection was achieved with LiteAblot Extend Chemiluminescent Substrate (EuroClone, Milan, Italy), and band density was quantified using the Invitrogen iBright 1500 Imaging Systems (Thermofisher Scientific, Waltham, Massachusetts, USA). For normalization, the optical density of each target band was compared to that of actin.

### 2.13 Wound healing assay

A standard scratch wound healing assay was employed to evaluate NHDF migration. NHDFs were seeded in triplicate at 300,000 cells/well in a 6-well plate, and a uniform wound was created in the confluent cell monolayer in each well using a sterile 200 µL micropipette tip. The dislodged cells and debris were then rinsed away with DMEM. The remaining adherent cells were treated with SnS at dilutions of 1:40, 1:60, and 1:80, prepared in FBS-free DMEM. A control group, maintained without SnS, was established in parallel. The wound sites were visualized at two time points (immediately after scratching, designated T0, and 24 h post-treatment) using an inverted light microscope (Leica DMi1, Leica Cambridge Ltd., Cambridge, UK) coupled with a camera and free Leica LAS EZ software. To quantify migration, the wound area at each time point was measured using Fiji ImageJ 1.54f software. The percentage of wound closure was calculated by normalizing the reduction in the wound area at 24 hours to the initial wound area measured at T0.

### 2.14 Statistical evaluation

Mean and standard deviation were calculated by R-Studio (v. 2025.05.01). The results of repeated readings were combined and displayed as mean ± SD. Differences at p < 0.05 were considered significant. The analysis was performed using the statistical software GraphPad Prism v.8.0 (San Diego, CA, USA) using one-way and two-way ANOVA followed by Dunnett’s test.

## 3. Results

### 3.1 Cell metabolic activity analysis

NHDF viability was measured by the Alamar blue assay after 24, 48 and 72 h of SnS treatment. After 24 h of treatment, no significant change in cell viability was detected, while after 48 h of treatment, SnS 1:40 significantly increased the cell viability, and after 72 h, SnS 1:60 significantly increased the cell viability with respect to CTRL (Figure 1).

**Figure 1:**
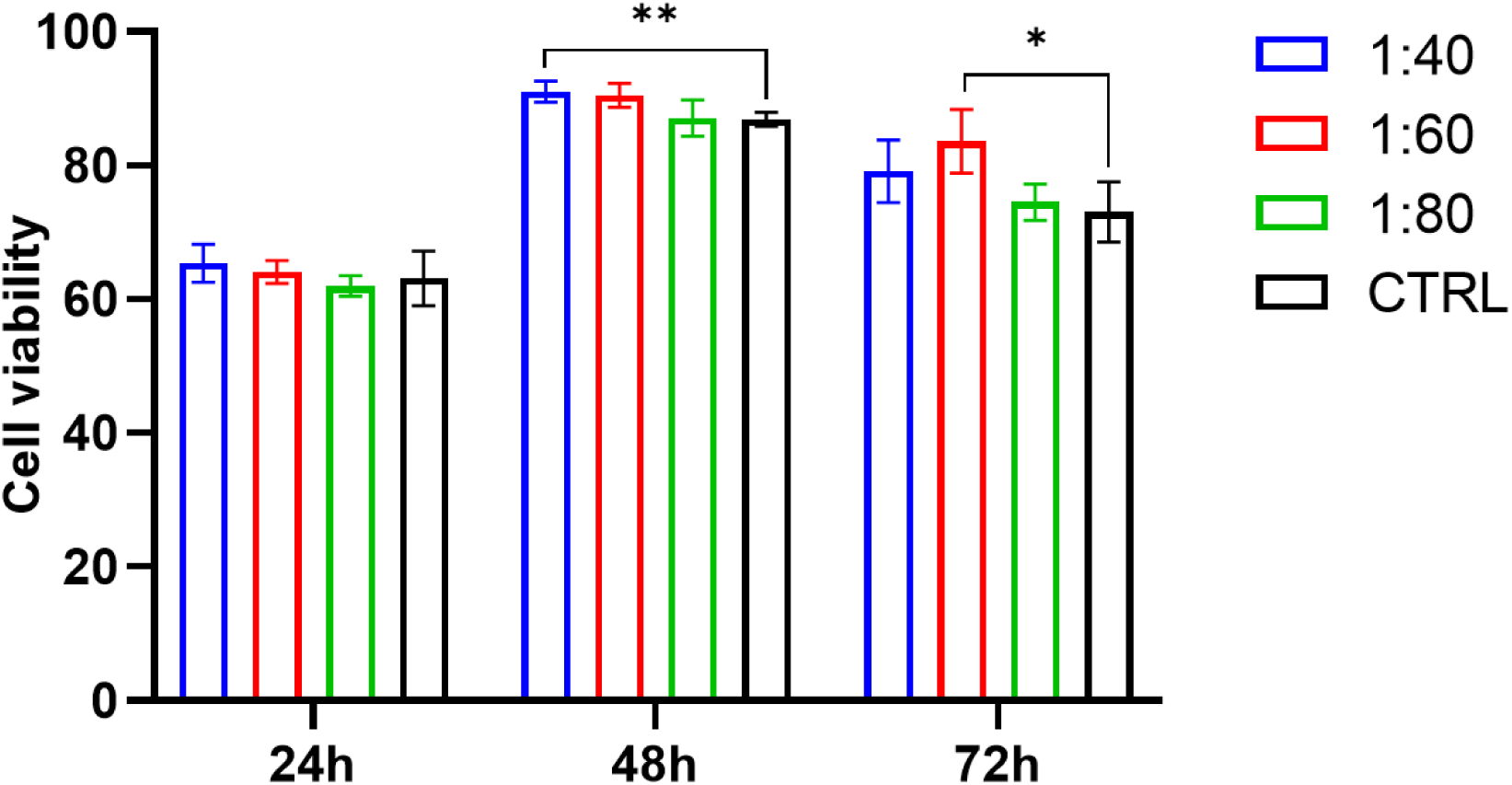
Cell metabolic activity assay using Alamar blue assay in NHDFs, treated with different concentrations of SnS (1:40, 1:60, 1:80) for 24, 48, and 72 h. The histogram represents Alamar blue reduction percentage equivalent to cell viability percentage; data shown are the mean (±SD) of three separate experiments. **p<0.001, *p<0.01.

### 3.2 Proteomics results

Being the major population of skin, NHDFs were chosen for further exploration of proteomics profiling after SnS treatment. The proteomics examination of NHDFs provided 5339 identified proteins (Supplementary Table 2). The number of identified proteins was relatively consistent among the groups; however, the sample group representing SnS 1:60 after 48 h treatment showed a slightly lower number of protein identifications (Supplementary Figure S1). A statistical analysis was employed to identify the DAPs (Supplementary Table 3). Venn diagram illustrates common and unique DAPs among proteins with increased or decreased abundance, respectively, and shows the number of DAPs obtained by comparing the groups to the CTRL. In the case of DAPs with increased amount, only 92 DAPs were common to all SnS concentrations (1:40, 1:60, 1:80) at 24 h, this number increased considerably to 261 after 48 hours, while in the case of DAPs with decreased amount, 119 proteins were common at 24 h changing to 263 after 48 h (Figure 2).

**Figure 2:**
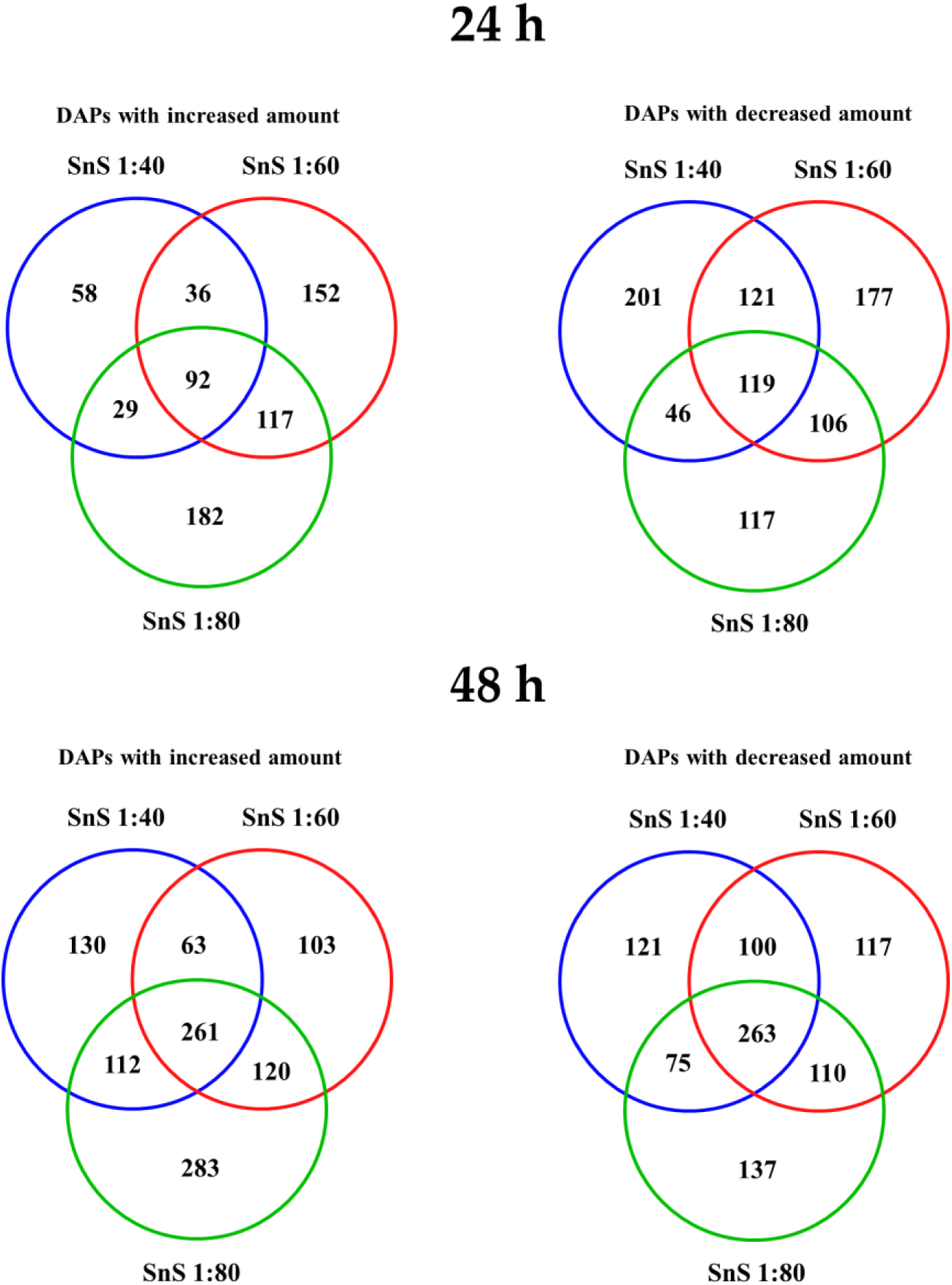
Venn diagrams showing the DAPs in each treatment group exposed to different dilutions of SnS after 24 and 48 h compared to the control.

#### 3.2.1 Network analysis

The protein-protein interaction (PPI) networks of DAPs and their first shell interactors were examined in string-db.org. For easier visualization we created combined networks for DAPs with either increased or decreased abundances, respectively, from all treatments: all SnS concentrations and both timepoints (Figure 3).

**Figure 3:**
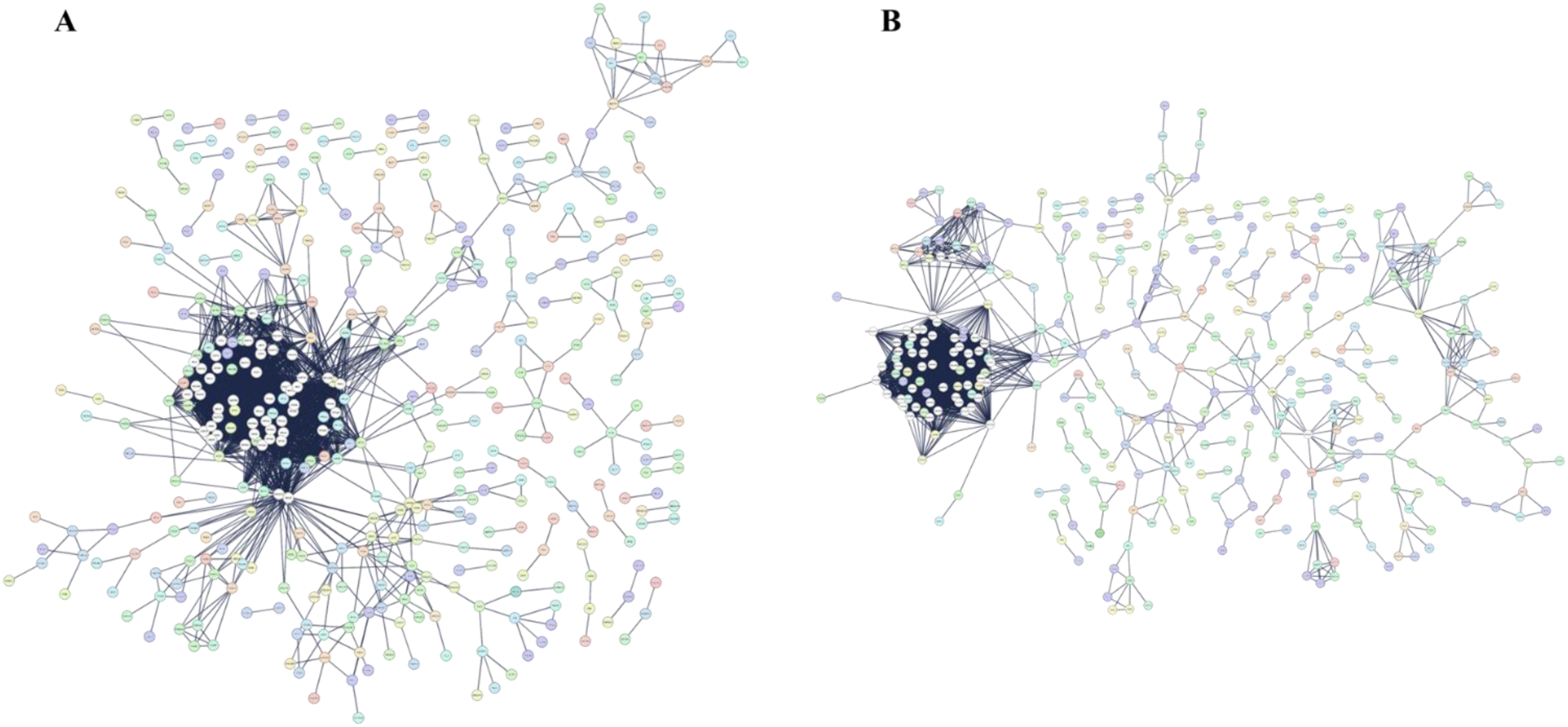
Network analysis of DAPs, with increased abundance (A) and decreased abundance (B). In the functional networks, circles represent DAPs and their first-shell interactors, while lines indicate PPI. The high resolution and magnifiable networks can be found in Supplementary Figures S2.

One highly connected cluster and several smaller clusters but different network architecture was observed in the case of networks containing DAPs with increased or decreased abundancy, respectively. The highly connected clusters correspond to functional modules and protein complexes that work together to perform specific biological processes.

Hub proteins, the proteins with the highest number of interactions in a network indicate proteins with higher importance. With network analysis we could identify the 10 most important hub proteins in our networks (Figure 4).

**Figure 4:**
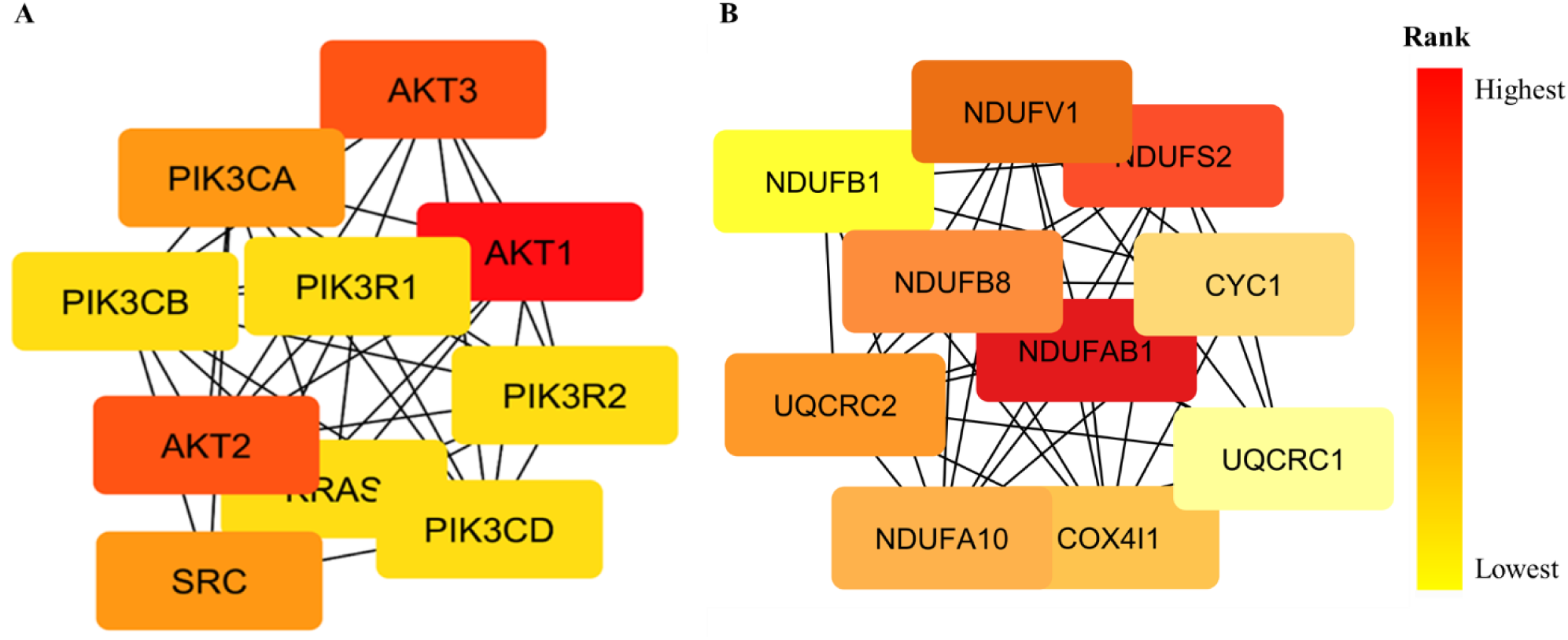
Top 10 hub proteins identified by CytoHubba Cytoscape plugin of DAPs, with increased abundance (A) and decreased abundance (B). The proteins are ranked highest to lowest based on the PPI by measuring the density of nodes.

Among the proteins increased or specifically appeared in the SnS treatment the AKT family members, also known as protein kinase B (PKB), a family of serine/threonine-specific protein kinases showed highest connectivity, followed by PI3K (Phosphoinositide 3-Kinase) family members, SRC (Proto-oncogene tyrosine-protein kinase Src) and KRAS (Figure 4A). All these proteins are involved in various cell signalling pathways that control cell growth, metabolism, cell survival, and related functions. The top 10 hub proteins identified in the case of PPI networks for DAPs with decreased amount following SnS treatment, a different picture could be observed: members of the NDUF protein family and other proteins involved in electron transport and mitochondrial energy production were the hubs (Figure 4B).

#### 3.2.2 Functional analysis

Functional analysis highlighted the most important functions of DAPs. According to the GO enrichment analysis the five most relevant biological functions (Figure 5) were the regulation of programmed cell death, followed by responses to oxidative stress, wound healing, positive regulation of cell migration and aging.

**Figure 5:**
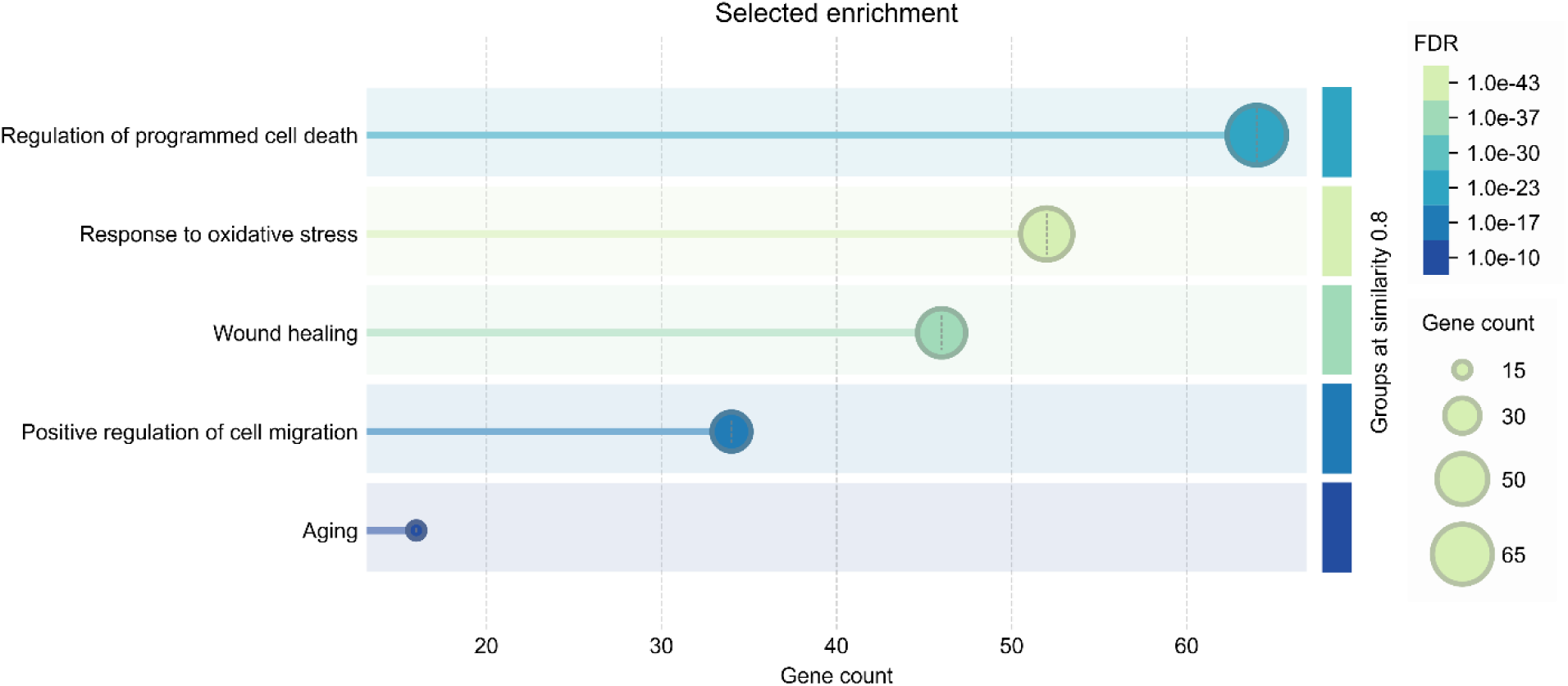
Functional enrichment analysis of DAPs. GO terms were sorted based on gene count; the FDR corrected q values are also indicated. The functional enrichment of each treatment group can be found in Supplementary Figures S2.

To corroborate the proteomic predictions at a functional level, complementary cellular assays were performed to assess the major biological processes and signaling pathways identified by network and enrichment analyses, including cell survival, migration, oxidative stress modulation, inflammatory signaling, and matrix remodeling.

### 3.3 Evaluation of programmed cell death occurrence

SnS effect on apoptosis and necrosis induction was analyzed through flow cytometry, in order to validate the proteomic data. After 24 h of SnS treatment a statistically significant increase of live cells percentage (Annexin V^-^/PI^-^) is detected comparing all the three SnS dilutions (1:40, 1:60 and 1:80) with the CTRL. The percentage of necrotic cells (Annexin V^-^/PI^+^) and early apoptotic cells (Annexin V^+^/PI^-^) after SnS administration appears comparable to those observed in the CTRL and, nevertheless, accounted for a negligible proportion (approximately 2%). The percentage of late apoptotic cells (Annexin V^+^/PI^+^) after treatment with 1:40, 1:60 and 1:80 is significantly lower compared to the percentage recorded in the CTRL group. After 48 h, an opposite pattern was observed: a slight, but statistically significant reduction (around 20%) in the percentage of live cells is detected after treatment with the three SnS dilution with respect to CTRL, with a major extent for 1:80 dilution. The amount of necrotic and early apoptotic cells remains unchanged compared with that observed after 24 h of treatment, while the percentage of late apoptotic cells is significantly increased after SnS exposure with respect to the CTRL, again with a major extent for 1:80 dilution (Figure 6).

**Figure 6:**
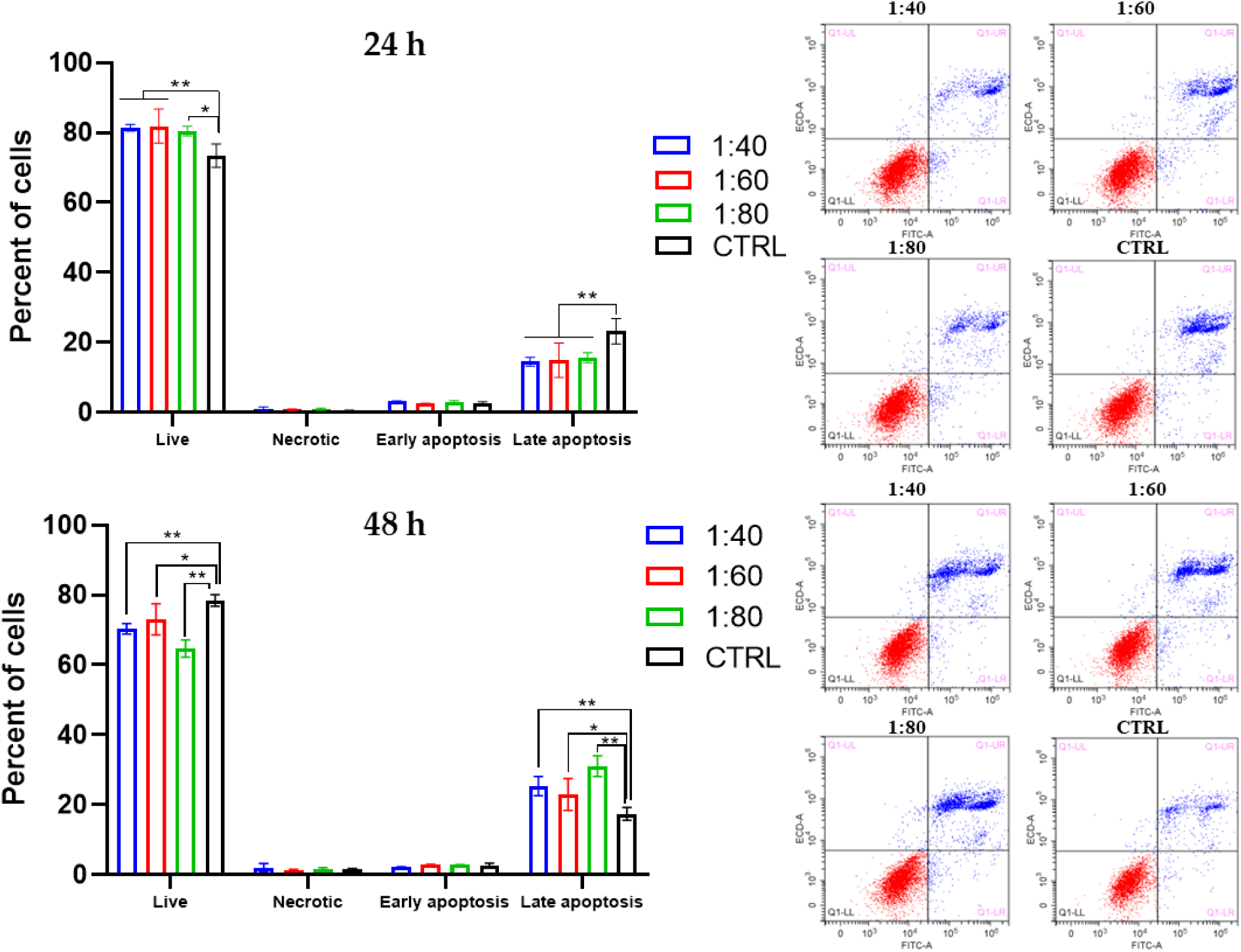
Flow cytometry analysis of apoptosis induction in NHDFs after 24 and 48 h of SnS (1:40, 1:60, 1:80) treatment. Percentages of Annexin/PI-stained cells analyzed by flow cytometry are shown. Histograms represent the percentage of total apoptotic cells (Annexin V^+^/PI^+^) reported as mean ± SD of three independent experiments. Representative dual-parameter fluorescence density dot plots are displayed close to the histograms. **p<0.001, *p<0.05.

The two main proteins that work antagonistically during the process of programmed cell death are the proapoptotic Bax and the antiapoptotic Bcl-2. To measure the effect of SnS treatment on programmed cell death in NHDFs the expression of Bax and Bcl-2 proteins were evaluated by western blot analysis. After 24 h of SnS treatment, Bax expression level in the SnS 1:40 and 1:80 treatment groups were significantly higher compared to control, while in 1:60 it was significantly lower; after 48 h of treatment Bax expression was significantly higher in SnS 1:40 and decreased significantly in SnS 1:60 and 1:80 compared to control (Figure 7A). Conversely, in the case of Bcl-2, all the SnS treated groups show significantly higher Bcl-2 expression than the control after 24 and 48 h of treatment (Figure 7B).

**Figure 7:**
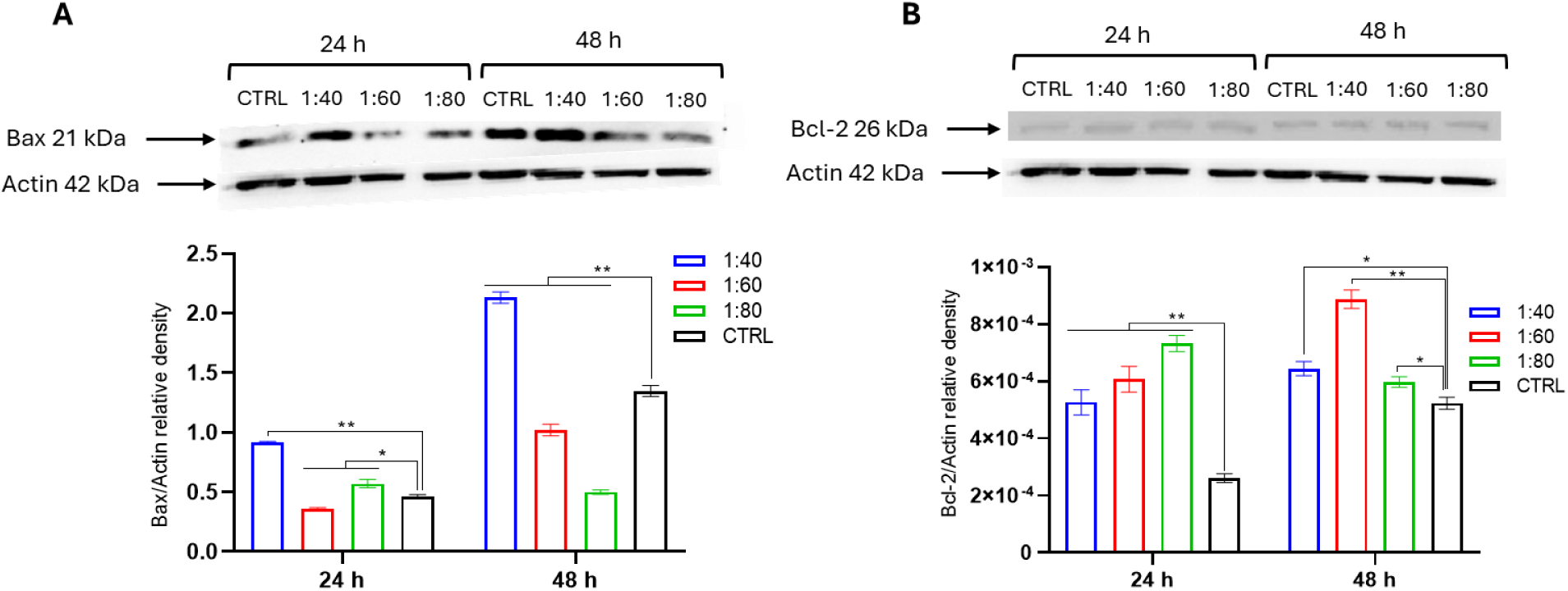
Western blot analysis of (A) Bax and (B) Bcl-2 in NHDFs exposed to different dilutions of SnS (1:40, 1:60, 1:80) and CTRL. The graph represents a densitometric analysis of protein expression levels normalized to actin expression levels. Data are presented as the mean ± SD of three independent experiments. **p<0.001, *p<0.05.

### 3.4 Antioxidant capacity assessment

The antioxidant capacity of NHDFs was quantified following a 24-hour treatment with various dilutions of SnS (1:40, 1:60, 1:80). A significant increase in antioxidant activity was observed in all SnS-treated groups compared to control, indicating that SnS treatment robustly enhances the free radical scavenging potential of dermal fibroblasts (Figure 8A).

**Figure 8.**
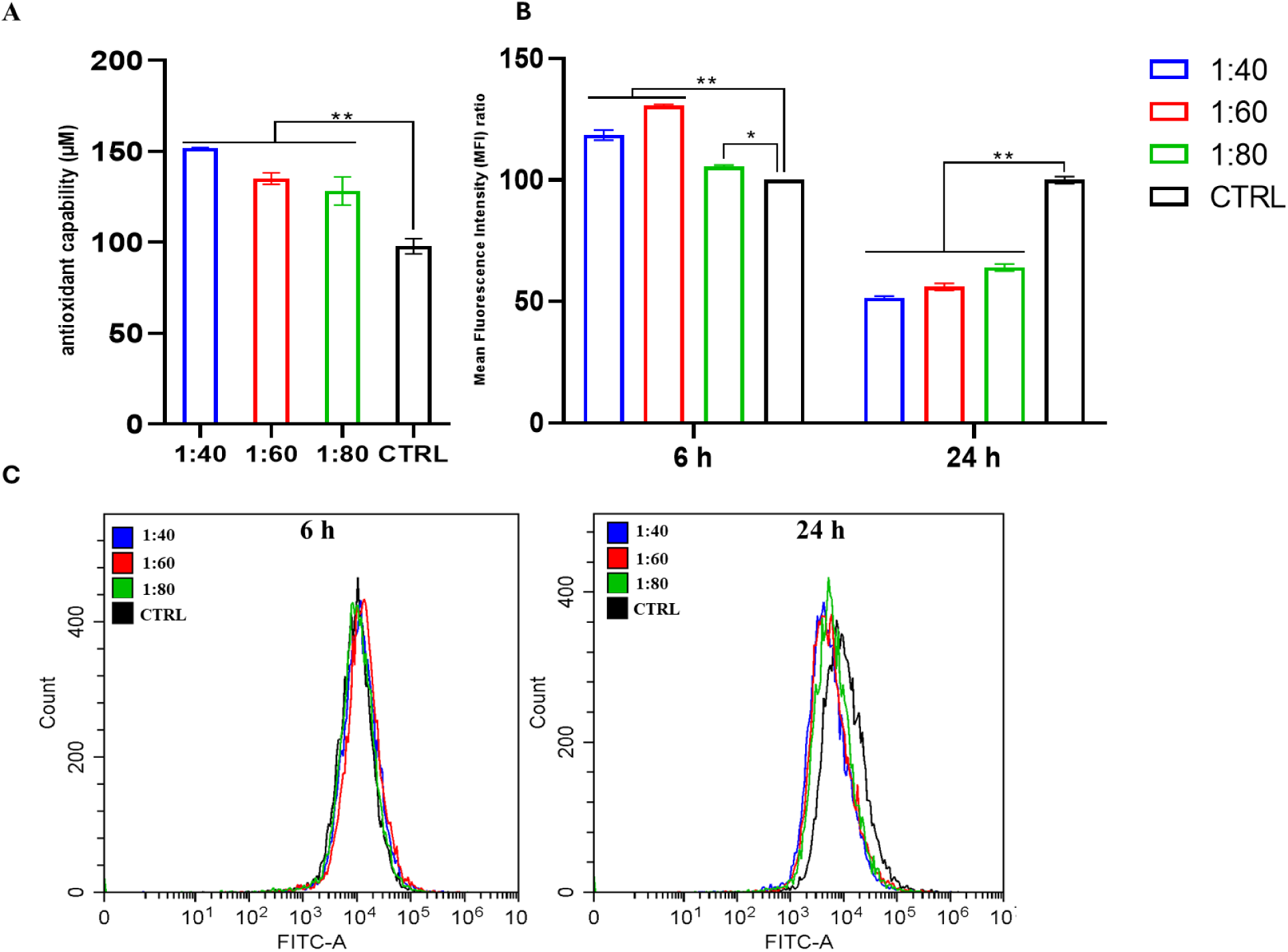
(**A**) Antioxidant capacity in NHDFs treated with 1:40, 1:60, and 1:80 dilutions of SnS for 24 h. Histograms represent median values ± SD of three independent experiments. Statistical analysis was performed using one-way ANOVA, followed by Dunnett’s multiple comparisons test. (**B**) Effects of SnS at dilutions of 1:40, 1:60, and 1:80 on intracellular ROS production in NHDFs. Intracellular ROS levels were assessed by flow cytometry after 6 and 24 h of treatment. The histogram represents the median fluorescence intensity (MFI), reported as fold change versus control (CTRL), and are expressed as mean ± SD of three independent experiments. (**C**) A representative overlay plot is shown. The Y-axis indicates MFI generated by the oxidation of H₂DCFDA, reflecting intracellular ROS production. **p < 0.001, *p < 0.05.

Intracellular ROS level in NHDFs was measured after 6 and 24 h of SnS (1:40, 1:60, 1:80) treatment by flow cytometry. After 6 h of treatment, the ROS release level was significantly higher in all the treatment groups compared to control, while after 24 h of treatment, SnS significantly reduced the ROS release level in all the treatment groups in a dose-dependent manner (Figure 8B and 8C).

### 3.5 Wound healing analysis

The effect of SnS on cell migration was evaluated using a scratch wound healing assay. Treatment with all SnS dilutions (1:40, 1:60, 1:80) resulted in a statistically significant enhancement of NHDF migration compared to the untreated control. Remarkably, all SnS-treated groups achieved complete wound closure (100%) within the 24-hour experimental period, demonstrating the potent pro-migratory activity of SnS (Figure 9).

**Figure 9.**
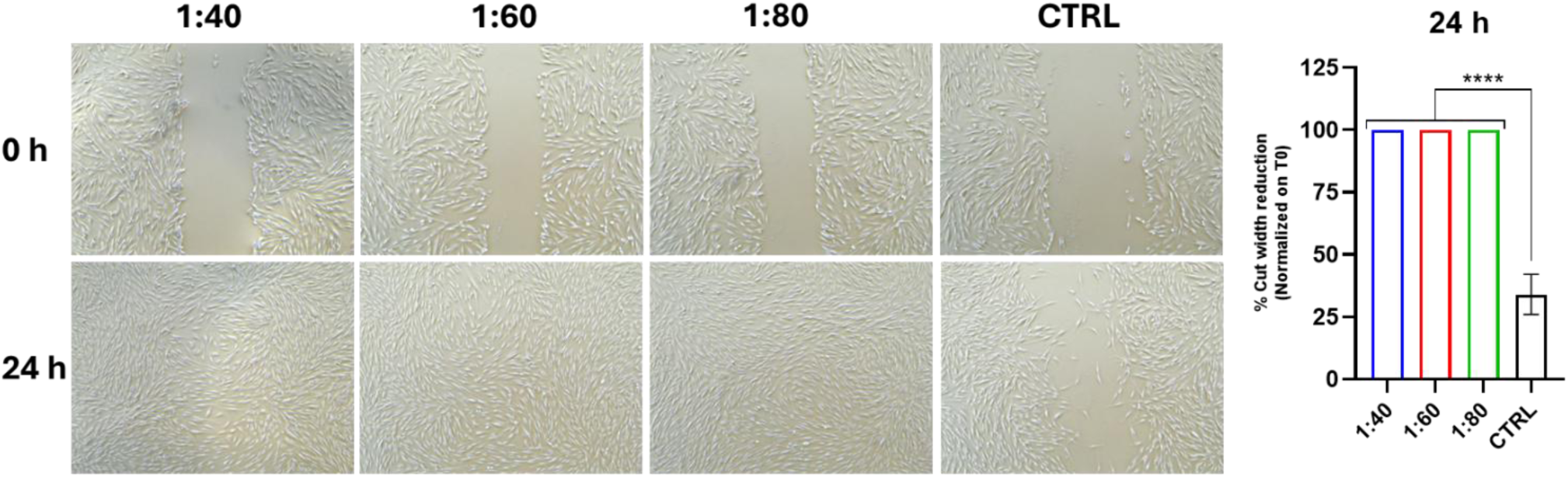
Wound healing assay in NHDFs after treatment with SnS at different concentrations. Representative images are reported, acquired at 5X magnification at 0 h and 24 h after treatment. Bar graph displays the percentage of cut width reduction after 24 h of treatment, normalized on the cut measurement at T0. ****p<0.0001 *vs* CTRL.

## 4. Discussion

Proteins serve as structural support, biochemical catalysts, hormones, enzymes, building blocks, and initiators of cellular death; on the other hand, proteins act as the machinery of cellular metabolism and communication [29]. To better understand the biological pathways involved in a physiological or pathological biological functions, the identification and/or measurement of the production and release of proteins represents a crucial step [30]. Omics approaches, including proteomics, not only simplified the ways to identify and quantify the proteins involved in biological functions, but also help to unfold the complex mixture of proteins [31].

SnS is proposed to significantly modulate the production and expression of key proteins involved in fundamental biological processes, such as cell proliferation, wound healing through cell migration, antioxidant defense and inflammatory responses [10, 15]. Thus, to better explore the SnS’s effect at the protein level, this study decided to employ an LC-MS-based proteomic approach and network analysis, further validated by various non-proteomic methodologies. Our method is an excellent example of data-driven approaches: the proteins identified and quantified by proteomics analysis, as well as their functional analyses, pinpointed some relevant pathways that were further tested by biochemical assays.

Dermal fibroblasts’ viability, assessed by the Alamar Blue assay, clearly evidenced that SnS is perfectly tolerated, even after 72 h of exposure, underlining the outcomes of our previous research on keratinocytes, HGFs and ECs [10, 15]. Considering the results of cell viability, we further deepen our findings on NHDFs at the proteomic level after SnS treatment, making this study the first ever to do so. The unique proteins obtained after applying filters from the DAP datasets were used for network and GO enrichment analysis confirming the role of SnS in five highly important biological functions: (i) regulation of programmed cell death, referred as apoptosis, (ii) reduction of oxidative stress, (iii) wound healing, (iv) cell migration, and (v) anti-aging. Many of these biological functions have already been confirmed by other researchers by non-proteomic approaches on different cell lines. Ho *et al*. [32] mentioned that *Achatina fulica* slime induced programmed cell death through the extrinsic apoptotic pathway in the triple-negative breast cancer cells, via activation of *Fas* pathway, while downregulating apoptosis in normal cells. Similarly, Cabibbo *et al*. [33] confirmed that SnS elevates the activities of antioxidant enzymes, such as glutathione and superoxide dismutase, and our group [15] reported the reduction in ROS release in HaCaT after SnS treatment, corroborating the role of SnS in counteracting the occurrence of oxidative stress. Functions such as wound healing, cell migration and anti-aging after SnS treatment are also well documented in previous studies of our group on SnS [10, 15]. The pathways related to apoptosis, cell migration, aging and wound healing, highlighted by the unbiased proteomics approach, could be validated by functional assays, providing further evidence for the beneficial effect of SnS on skin functions.

Proteins with the highest number of connections in the network, the so-called hub proteins, can give information on the importance of a protein in the network [34]. When the protein abundance changes, the possibility of interactions is changed, which can affect the overall network architecture and the hub proteins [35]. The network of proteins whose abundance decreased upon SnS treatment showed a highly connected cluster of mostly NDUF proteins and several smaller clusters. As expected, based on the network architecture, the hub proteins predominantly belonged to the NDUF protein family, which is involved in electron transport and mitochondrial energy production, likely reflecting reduced cellular energy metabolism. In contrast, the network of proteins whose abundance increased had a different architecture: a highly connected cluster containing various proteins and the hub proteins were members of AKT, PI3K, SRC and KRAS protein families. In detail, AKT plays a key role in the PI3K/Akt signaling pathway, promoting cell proliferation and preventing apoptosis; PI3Ks control vital cellular functions by key PI3K/Akt/mTOR signaling pathways and regulate cell growth, metabolism, and survival [36]., while the SRC protein acts as a crucial signaling molecule regulating cell growth, proliferation, migration, adhesion, and survival [37]. Finally, the KRAS protein is a key part of the RAS/MAPK signaling pathway that acts like an “on-off” switch for cell growth and division; in healthy cells, this cycle controls growth, differentiation, and cell survival [38]. Collectively, the enrichment of AKT, PI3K, SRC, and KRAS family members as hub proteins indicates that SnS-driven proteomic changes converge on central signaling nodes that coordinate multiple repair-related processes rather than acting on isolated pathways. These hub proteins integrate survival, migratory, and stress-response signals, thereby synchronizing apoptosis regulation, antioxidant capability and proliferative responses required for effective wound healing. Their high connectivity suggests that SnS modulates upstream regulatory points capable of orchestrating broad cellular programs, enabling a coordinated balance between cell survival, controlled turnover, and migration. This network-level regulation provides a mechanistic basis for the observed enrichment of biological functions such as apoptosis modulation, wound healing, and cell migration.

The regulation of programmed cell death, more specifically apoptosis, was cross-verified by Annexin/PI staining, by flow cytometry, with an additional investigation of Bax and Bcl-2 expression, two main regulators of the apoptotic process. The increase of living cells after 24 h of SnS treatment, along with the reduction of apoptotic cells compared to control (no apoptosis induction), combined with the opposite situation recorded after 48 h, in which a reduction of SnS-treated living cells and an increase of apoptotic events occurs, suggested that SnS can act over time triggering a turnover and thus leading to a beneficial cell renewal. Indeed, from 24 h, the level of Bax increases in the 1:40 and 1:80 samples, likey triggering an activation of the intrinsic apoptotic cascade, thus supporting the increase of apoptotic events recorded by Annexin/PI after 48 h. The 1:60 dilution, even inducing the same Annexin/PI result obtained with 1:40 and 1:80, does not induce the same Bax higher level: considering that the apoptotic process is orchestrated by different pathways and that Bax is only one of the actors involved [39, 40], it can be argued that in the 1:60 dilution other molecules, such as those related to the extrinsic apoptotic pathway, are involved to obtain the same result induced by 1:40 and 1:80. Thus, the increment of apoptotic cells can be explained admitting an induction of cellular turnover for two main reasons: I) the reduction of living cells and the increase of apoptotic ones, recorded by Annexin/PI, is low, approximately 15-20%, therefore is not a massive effect; II) the Alamar blue data, at 48 and 72 h, indicate that cells remain healthy and viable after prolonged treatment. This hypothesis is corroborated by the reduced levels of Bax after 48 h, suggesting a transitory apoptotic event, correctly balanced by the activity of anti-apoptotic proteins, such as Bcl-2, in order to prevent a marked apoptosis induction. In fact, it can be evidenced that Bax and Bcl-2 show the same trend: the higher Bax levels are paralleled by increased Bcl-2 levels, indicating that Bcl-2 acts to counterbalance the pro-apoptotic effect of Bax, as elsewhere confirmed by Qian *et al*. [41] which state that Bcl-2 is able to inhibit apoptosis by forming a heterodimer with Bax and to ensure cell survival by regulating the Ca^+2^ concentration and antioxidant effect. Additionally, it can also prevent apoptosis by inhibiting the activities of caspase-9, 3, 6, and 7.

Oxidative stress is a central regulator of fibroblast function, where controlled ROS release acts as critical signaling mediators in wound healing, while excessive or sustained ROS accumulation impairs regeneration and induces cellular damage [42]. In this study, SnS finely modulated intracellular redox homeostasis, inducing a biphasic ROS response characterized by a transient early increase at 6 h followed by a pronounced reduction at 24 h across all tested dilutions. This pattern reflects a hormetic redox effect, where an initial oxidative signal primes endogenous defense mechanisms, ultimately enhancing antioxidant capacity and cellular resilience, making these results coherent with previous findings [15, 22, 33]. Importantly, this redox modulation is supported by proteomic evidence: Gene Ontology analysis revealed, after SnS treatment, an enrichment of proteins involved in oxidative stress responses, indicating coordinated remodeling of redox-regulatory networks. In fact, it has already been reported by Teng *et al.* [43], that the activation of the PI3K/Akt signaling pathway confers protection against oxidative stress-induced apoptosis in melanocytes. In particular, PI3K/Akt activation is associated with increased expression of anti-apoptotic Bcl-2 and concomitant downregulation of pro-apoptotic Bax and caspases-3 and -9 [32, 44]. These findings align with previous studies showing that SnS is also able to enhance key antioxidant enzymes, including glutathione peroxidase and superoxide dismutase, thereby reinforcing intrinsic detoxification systems [32, 45].

Functionally, the ROS dynamics induced by SnS were accompanied by regulated apoptotic signaling, with controlled Bax modulation and elevated Bcl-2 expression, supporting cell survival over oxidative stress–driven death. Simultaneously, the reduction in ROS at 24 h, coupled with enhanced antioxidant capacity, likely establishes a permissive environment for fibroblast migration, which may be mediated through redox-sensitive activation of the PI3K/AKT signaling pathway, promoting cytoskeletal remodeling and pro-migratory programs [43]. This dual effect, attenuating oxidative stress while sustaining pro-migratory signaling, positions SnS as a redox-modulating biomaterial capable of orchestrating a dynamic balance between ROS signaling, antioxidant protection, and regenerative activity. These mechanistic insights, integrated with the proteomic reprogramming observed, provide a robust rationale for the pro-healing and anti-aging effects of SnS in human dermal fibroblasts.

The most striking finding is the potent pro-migratory activity of SnS that regulates the wound healing and aging process. Thus, to confirm the proteomic data, the scratch assay results were assessed revealing that all tested dilutions of SnS significantly accelerated wound closure, achieving 100% gap closure within 24 h, a stark contrast to the control. This suggests that SnS contains bioactive compounds that powerfully stimulate the cellular migration essential for wound repair, already witnessed by previous findings [46–48]. It has been in fact demonstrated that a superficial scar in albino rats treated with SnS achieved 64% of healing after 14 days, compared to a standard Madecassol ointment, which took 21 days [47]. Similarly, Gubitosa *et al*. mentioned that gold nanoparticles of SnS significantly increase the urokinase-type plasminogen activator receptor (uPAR), essential for keratinocyte adhesion, spreading, and migration [48]. On the ither hand, we noticed that SnS enhanced the cellular antioxidant capacity, indicating a role in bolstering cellular defense against oxidative stress. The profound reprogramming of the cellular proteome observed, suggests that SnS may induce a controlled stress response that redirects energy from proliferation to migration and defense mechanisms. This action, promoting migration while enhancing antioxidant activity, positions SnS as a promising agent for supporting wound healing, where rapid tissue closure and protection from ROS are critical passages.

## 5. Conclusion

The present study explores and validates SnS-induced proteome-wide alterations in NHDFs, providing mechanistic insight into its biological activity. Network-based and functional enrichment analyses demonstrate that SnS modulates interconnected protein clusters governing key cellular processes involved in tissue repair. Notably, SnS treatment upregulated proteins associated with cell viability and migratory capacity, both of which are critical for efficient wound closure. Concurrently, proteins related to oxidative stress responses and programmed cellular turnover were modulated, suggesting that SnS favours adaptive, protective stress signaling rather than detrimental cellular damage. The proteomic signature reflects a highly coordinated, enhanced cytoprotective defense system that preserves cellular integrity while promoting regenerative processes.

Overall, SnS does not merely function as a growth-promoting agent but acts as a sophisticated biological modulator. Integrated proteomics and functional analyses confirm its multifaceted role in regulating cell viability, apoptosis, oxidative stress mitigation, and wound healing. These findings highlight the therapeutic potential of SnS and warrant further investigation into the specific signaling pathways and key regulatory proteins underlying its biological effects.

## Author Contributions

Conceptualization SZ, EC; Methodology: MR, AR, MB, ZD, AP; Software: MR, AR, MB, AP; Validation: SZ, EC; Formal Analysis: AR, MB; Investigation: MR, AR, MB; Resources: SZ, EC, ZD; Data Curation: MR, AR, AP; Writing—Original Draft Preparation: MR, AR, MB, AP; Writing—Review & Editing: SZ, EC, ZD; Visualization: AC, SZ; Supervision: SZ, EC, Project Administration: SZ; Funding Acquisition: MR.

## Acknowledgments

This publication was produced during MR’s attendance at the PhD program in Biomolecular and Pharmaceutical Sciences at “G. d’Annunzio” University of Chieti-Pescara, Italy, Cycle XXXVIII, with the support of a scholarship financed by the Ministerial Decree no. 351 of 9th April 2022, based on the National Recovery and Resilience Plan (NRRP) - funded by the European Union - NextGenerationEU - Mission 4 “Education and Research”, Component 1 “Enhancement of the offer of educational services: from nurseries to universities” - Investment 3.4 “Advanced teaching and university skills”. We thank the staff of the Proteomics Core Facility, Department of Biochemistry and Molecular Biology, Faculty of Medicine, University of Debrecen, Debrecen, Hungary for providing the research facilities and for hosting MR during his attendance of PhD program. We thank the technical help for Mabuse Moagi Gontse, Mahshid Mobalegh Naseri, Mansi Jain, Julia Kökényesiné Csáki and Dr. Gergő Kalló. The results presented in the current paper were obtained thanks to the collaboration between University “G. d’Annunzio” Chieti-Pescara and Lumacheria Italiana S.r.l. (Cherasco, Italy) in accordance with the agreement signed on 4 February 2025.

